# Estrogen receptor β exerts tumor suppressive effects in prostate cancer through repression of androgen receptor activity

**DOI:** 10.1101/850180

**Authors:** Surendra Chaurasiya, Scott Widmann, Cindy Botero, Chin-Yo Lin, Anders M. Strom, Jan-Åke Gustafsson

## Abstract

Estrogen receptor β (ERβ) was first identified in the rodent prostate and is abundantly expressed in human and rodent prostate epithelium, stroma, immune cells and endothelium of the blood vessels. In the prostates of mice with inactivated ERβ, mutant phenotypes include epithelial hyperplasia and increased expression of androgen receptor (AR)-regulated genes, most of which are also upregulated in prostate cancer (PCa). ERβ is expressed in both basal and luminal cells in the prostate while AR is expressed in luminal but not in the basal cell layer which harbors the prostate stem cells. To investigate the mechanisms of action of ERβ and its potential cross-talk with AR, we used RNA-seq to study the effects of estradiol or the synthetic ligand, LY3201, in AR-positive LNCaP PCa cells which had been engineered to express ERβ. Transcriptomic analysis indicated relatively few changes in gene expression with ERβ overexpression, but robust responses following ligand treatments. There is significant overlap of responsive genes between the two ligands, as well as ligand-specific alterations. Gene set analysis of down-regulated genes identified an enrichment of androgen-responsive genes, such as FKBP5, CAMKK2, and TBC1D4. Consistently, AR transcript, protein levels, and transcriptional activity were down-regulated following ERβ activation. In agreement with this, we find that the phosphorylation of the CAMKK2 target, AMPK, was repressed by ligand-activated ERβ. Down-regulation of TBC1D4, a major regulator of glucose uptake in prostate, indicates that ERβ is changing glucose metabolism in the prostate. These findings suggest that ERβ-mediated signaling pathways are involved in the negative regulation of AR expression and activity, thus supporting a tumor suppressive role for ERβ in PCa.

## Introduction

The prostate is an androgen-responsive organ which is controlled by the androgen receptor (AR) both under normal physiological conditions and in malignancy. AR plays an important role in PCa as a strong driver of proliferation, and as such is the primary target for treatment of PCa [1]. Recently it was shown that androgen signaling is essential for PCa tumorigenesis from prostatic basal cells[2]. In addition to the 2.5 million people living with PCa, 250,000 new cases are diagnosed with 28,000 men dying from PCa every year in the United States alone. Treatment strategies differ depending on factors such as the stage of the disease, the age and physical status of the patient, and what treatments patients have previously received. PCa is a very indolent cancer and can initially be treated with surgery or androgen deprivation therapy (ADT), however, ADT often leads to emergence of a more aggressive castrate resistant metastatic cancer (CRPCa) [3]. CAMKK2 has been identified as a key downstream target of AR in coordinating PCa cell growth, survival, and migration. In addition, this protein kinase affects bone remodeling and macrophage function, and is emerging as a therapeutic target downstream of AR for controlling metastatic PCa and preventing ADT-induced bone loss[4].

In addition to AR, the prostate also expresses estrogen receptor β (ERβ), which is expressed in both basal and luminal cells [5]. ERβ, a member of the nuclear receptor family, was discovered in 1996 [6]. This receptor has been shown to act as a tumor suppressor in several types of cancer [7-9]. In addition, when ERβ is deleted from the mouse genome, epithelial hyperplasia and upregulation of AR-regulated genes in the prostate occur[10]. ERβ is ligand-activated, which like ERα, can be activated by estradiol. However, in the prostate the more abundant ERβ ligand is not 17β-estradiol, but 17β-Adiol [11]. The basal cells in the prostate which are thought to harbor the prostatic stem cells, are AR-negative. Deletion of ERβ in the prostate gives rise to hyper proliferation of the epithelial layer and increases the proportion of intermediary luminal cells (luminal cells which are not fully differentiated) [10]. ERβ has previously been shown to reduce AR activity in the prostate by increasing the expression of the co-repressor DACH1/2 [12] and decreasing the AR driver RORc. In addition to DACH1/2, the co-repressor NCoR2/SMRT has also been shown to affect the activity of AR [13]. The overall effect of ERβ on gene networks in PCa cells has not been reported. The present study describes the first transcriptomic analysis of ERβ activation in AR-positive PCa cells and reveals a key role for ERβ in regulating AR expression and activity in PCa.

## Materials and Methods

### Reagents and cell culture

The LNCaP cell line was obtained from the American Type Culture Collection (ATCC). LNCaP cells were maintained in RPMI-1640 (Invitrogen Inc., Carlsbad, CA) medium supplemented with 10% fetal bovine serum (FBS) (Sigma, St. Louis, MO), and Antibiotic-Antimycotic (Invitrogen Inc., Carlsbad, CA). All experiments used cells below passage 30. R1881, 17β-estradiol, 4OH-tamoxifen, DHT (Sigma St. Louis, MO).

### LNCaP Cell lines expressing control virus or ERβ containing virus

LNCaP cells were infected with the lentivirus Lenti6-TOPO-V5-D, empty or containing cDNA for human ERβ at 2 m.o.i (multiples of infection). The control in all experiments is cells infected with empty virus vector.

### Protein extract preparation

To prepare whole-cell extracts, cells were washed twice with PBS, lysed in 10 times packed cell volume of lysis buffer [0.1% Nonidet P-40, 250 mM KCl, 5 mM Hepes, pH 7.9, 10% (vol/vol) glycerol, 4 mM NaF, 4 mM sodium orthovanadate, 0.2 mM EDTA, 0.2 mM EGTA, 1 mM dithiothreitol, 1 mM phenylmethylsulfonyl fluoride, protease inhibitor cocktail, PhosStop (Roche, Indianapolis, IN)] for 15 minutes on ice and then centrifuged at 14,000 x g for 10 minutes.

### Western blotting

Thirty micrograms of protein were loaded on an SDS-PAGE 10% Bis-Tris gel with Tris running buffer and transferred to a nitrocellulose membrane after electrophoretic separation. Membranes were blocked with 5% non-fat powdered milk in 0.1% TBST buffer and probed with anti-AR (sc-816), HSD11B2 (sc-365529), GAPDH-HRP (sc-47724), p57 KIP2 (sc-1037), E-cadherin (sc-7870), Cytokeratin 19 (sc-6278), TGFBR3 (sc-74511) (Santa Cruz Biotechnology,Inc., Santa Cruz, CA), Jagged 1 (70109T), RhoB (63876S), AMPK (2532S), pAMPK 2535S, FKBP5 122105 (Cell Signaling Technology Danvers, MA), CAMKK2 (H00010645-M01) (Abnova Taipei, Taiwan), p63 (ab124762), Ki67 (ab15580) (Abcam Cambridge, MA), β-Actin (A1978) (Sigma Millipore Sigma, St. Louis, MO). Primary antibodies were used at 1:200–1000 dilutions, and secondary antibody was used at 1:10,000. The western blot experiments were minimally repeated two times.

### RNA extraction and real-time PCR

RNA extraction was performed with Qiagen mRNA extraction kit according to standard protocol. cDNA was synthesized from 1μg of total RNA with First Strand System according to standard protocol (Invitrogen Inc. NY). Real-time PCR was performed with SYBR Green I dye master mix (Applied Biosystems Foster City, CA). Primers (Integrated DNA Technologies, Inc. Coralville, IA) were: GAPDH; F, 5-TGACAACTTTGGTATCGTGGAAGG-3 and R, 5-AGGCAGGGATGATGTTCTGGAGAG-3 (reference gene); AR; F, 5’-TCACCAAGCTCCTGGACTCC-3’ R, 5’-CGCTCACCATGTGTGACTTGA 3’; FKBP5; F, 5’-ATTGGAGCAGGCTGCCATTGTC -3’, R, 5’-CCGCATGTATTTGCCTCCCTTG-3’, CAMKK2; F, 5’-TCCAGACCAGCCCCGACATAG-3’, R, 5’-CAGGGGTGCAGCTTGATTTC-3’ 7500 Fast Real-Time PCR System (Applied Biosystems) using optimized conditions for SYBRGreen I dye system: 50 C for 2 minutes, 95 C for 10 minutes, followed by 40–50 cycles at 95 C for 15 seconds and 60 C for 50 seconds. Optimum primer concentration was determined in preliminary experiments, and amplification specificity confirmed by dissociation curve. The variance between the groups that are being statistically compared is similar.

### RNA-sequencing

Poly(A) mRNA was isolated using NEBNext poly(A) mRNA Magnetic Isolation Module. Libraries were prepared using NEBNext Ultra II RNA Library Prep Kit for Illumina. Sequencing was performed on NovaSeq 6000 with 150 bp paired-end reads. Treatments include three independent replicates.

### Transcriptome Analysis

Reads were aligned to reference genome (GRCh38) indexes using STAR^20^ (v2.5).. HTSeq^21^ (v0.6.1) was used for mapped gene count quantification. Differential expression analysis was performed using DESeq2^19^ (1.24.0). The resulting P-values were adjusted using the Benjamini and Hochberg’s method. Genes with an adjusted P-value <0.05 found by DESeq2 were assigned as differentially expressed. Venn diagrams and heat map were prepared in R. Gene set enrichment analysis (GSEA) ^17, 18^ was performed using rankings based on the test statistic from differential expression analysis and the hallmarks gene set (h.all.v7.0.symbols.gmt).

Gene Ontology (GO) enrichment analysis of differentially expressed genes was implemented by the cluster Profiler R package, in which gene length bias was corrected. GO terms with corrected P-values less than 0.05 were considered significantly enriched by differentially expressed genes. The RNA-seq data is available in NCBI’s Gene Expression Omnibus through accession number GSEXXXXXX.

## Results

### The transcriptomic effects of ERβ in the AR-positive cell line LNCaP

To determine the functions of ERβ in AR-positive PCa, we used RNA-seq to compare the transcriptomes of ERβ over-expressing and non-expressing LNCAP cells treated with vehicle (DMSO), and ERβ ligands estradiol (E2) and LY3201. Differential expression analysis consisted of comparing over-vs non-expressing ERβ within each treatment regime. DMSO-treatment treatment had very little effect on gene expression (Fig. 1A (heat map)). E2 and LY3201 treatments elicited changes in 4185 and 3456 differentially expressed genes (adjusted p-value < 0.05), respectively. The proportion of upregulated (60%) and downregulated (40%) genes between the two ligands was similar (Table 1). With the upregulated genes there was a strong overlap between the two treatments (63%). However, there was only a modest overlap in the downregulated genes show (46%). Ligand-dependent differentially expressed genes were less prevalent in LY3201 treated cells (Fig. 1B).

**Table 1.** Summary of numbers of differentially expressed genes identified in RNAseq study.

| Treatment | Differentially Expressed Genes* |  |  |
| --- | --- | --- | --- |
|  | Up-regulated | Down-regulated | Total |
| Vehicle (DMSO) | 29 | 14 | 43 |
| E2 | 2354 | 1831 | 4185 |
| LY | 2062 | 1394 | 3456 |

\*Defined as those with FDR-adjusted $p < 0.05$ .

**Figure 1.**
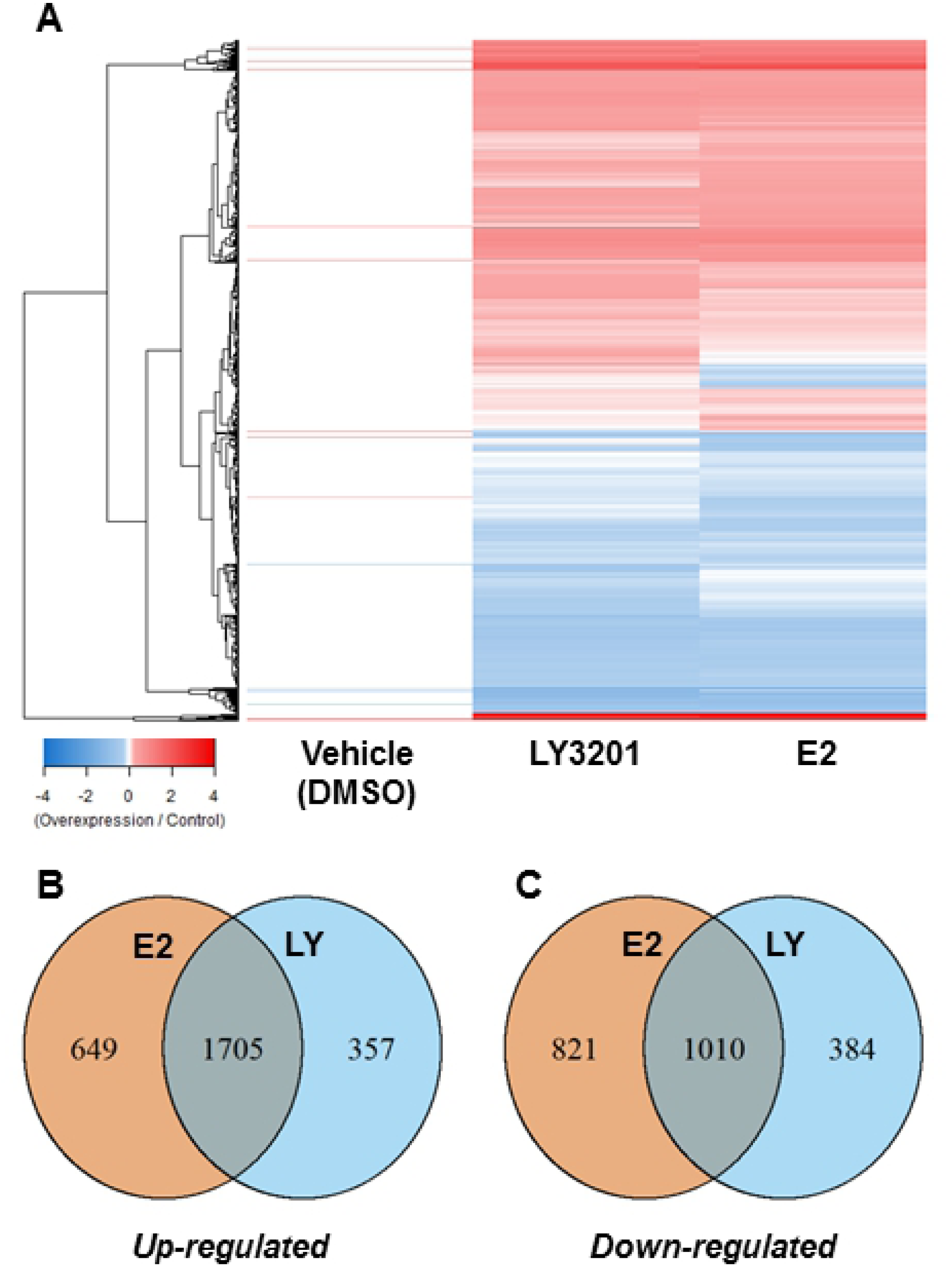
RNAseq analysis identified differentially expressed genes which respond to ERβ activation. A, Gene expression profiles of responsive genes in cells treated with vehicle, E2, or LY3021 are presented in a heatmap of log2-transformed fold-change values. Hierarchical clustering of expression profiles and resulting dendrogram group genes based on their similarities in responses to ligand treatment. B, Venn diagrams show the proportion of responsive genes which are common and specific to each of the two ERβ ligands used in the study.

Pre-ranked gene set enrichment analysis (GSEA) was used to identify pathways and mechanisms relevant to ERβ expression and activation in PCa. LY3201 was chosen for enrichment analysis. The most positively enriched set in the LY3201-treated cells were genes responsive to estrogen; other highly enriched sets contained genes involved in MTORC1 signaling and the unfolded protein response. GSEA also revealed downregulated genes involved in several pathways associated with cancer hallmarks such as hypoxia and glycolysis (Table 2). The most negatively enriched set contained genes involved in response to androgens a central driver in androgen-sensitive PCA (see Figure 2A). These androgen-responsive genes were investigated further due to the well-established importance of androgen signaling in prostate cancer. For full list of regulated genes see supplementary table S1.

**Table 2.** Summary of top hits from Gene Set Enrichment Analysis (GSEA) of ERβ ligand-responsive differentially expressed genes identified in RNA-seq study.

| Gene Set | NES | FDR q-value* |
| --- | --- | --- |
| *Positive Enrichment Score* |  |  |
| HALLMARK_ESTROGEN_RESPONSE_EARLY | 7.2 | 0 |
| HALLMARK_ESTROGEN_RESPONSE_LATE | 6.0 | 0 |
| HALLMARK_MTORC1_SIGNALING | 4.3 | 0 |
| HALLMARK_UNFOLDED_PROTEIN_RESPONSE | 3.7 | 0 |
| HALLMARK_E2F_TARGETS | 3.7 | 0 |
| HALLMARK_G2M_CHECKPOINT | 3.2 | 0 |
| HALLMARK_PROTEIN_SECRETION | 3.2 | 0 |
| HALLMARK_UV_RESPONSE_UP | 3.1 | 0 |
| HALLMARK_OXIDATIVE_PHOSPHORYLATION | 2.7 | 0 |
| HALLMARK_MITOTIC_SPINDLE | 2.5 | 3.9E-04 |
| *Negative Enrichment Score* |  |  |
| HALLMARK_ANDROGEN_RESPONSE | -3.4 | 0 |
| HALLMARK_HYPOXIA | -2.5 | 9.5E-04 |
| HALLMARK_P53_PATHWAY | -2.4 | 0.002 |
| HALLMARK_GLYCOLYSIS | -2.2 | 0.007 |
| HALLMARK_EPITHELIAL_MESENCHYMAL_TRANSITION | -2.0 | 0.01 |
| HALLMARK_KRAS_SIGNALING_UP | -2.0 | 0.02 |
| HALLMARK_UV_RESPONSE_DN | -2.0 | 0.02 |
| HALLMARK_WNT_BETA_CATENIN_SIGNALING | -1.9 | 0.02 |

\*FDR $q < 0.05$

**Figure 2.**
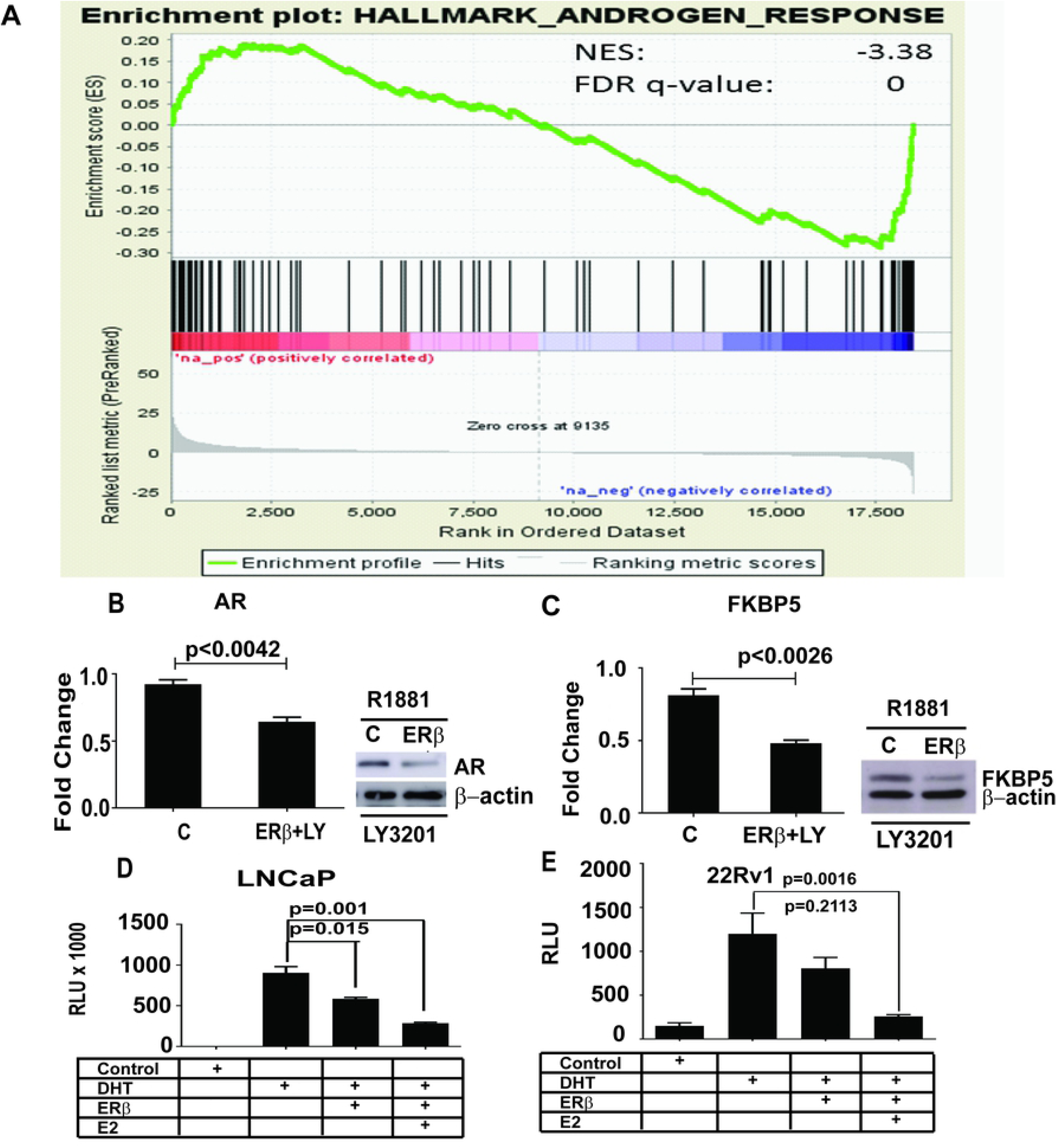
ERβ activation inhibit AR expression and transcriptional activity. A, Gene Set Enrichment Analysis (GSEA) revealed an enrichment of genes involved in androgen response down-regulated by ERβ activation (negative enrichment (NES) score). B, Transcript (graphs) and protein (western images) levels of AR and AR target gene FKBP5 were reduced by ERβ expression and treatment with LY3021. C, AR transcriptional activity was reduced by ERβ expression and activation in reporter gene assays in AR-positive LNCaP and 22Rv1 prostate cancer cells.

Table 1 showing differentially expressed genes up-regulated and down-regulated after treatment with E2 or LY3201

Table 2 showing hallmarks from Gene Set Enrichment Analysis (GSEA) positive and negative enrichment scores. NES Normalized Enrichment Score and FDR q-value False Discovery Rate.

### ERβ reduces the AR activity in LNCaP cells

Since AR-responsive genes were affected by ERβ activation (see Figure 2A), we investigated whether expression and/ or activity of AR was changed by expression of ERβ as has been proposed by several studies based on ERβ function in the mouse [11, 12]. We found that AR mRNA and protein were down-regulated by both estradiol and LY3201 in ERβ-expressing LNCaP cells (see Figure 2B). In addition, the established AR-regulated gene FKBP5 was also down-regulated (see 2igure 2C). Other established AR targets *(NDRG1, B2M, SORD, TPD52, DHCR24, ADAMTS1, NKX3A, RAB4, ANKH, TSC22, MAF [14]) were similarly down-regulated by ERβ ligand treatments according to the RNA-seq data (See Table 3).

**Table 3.**
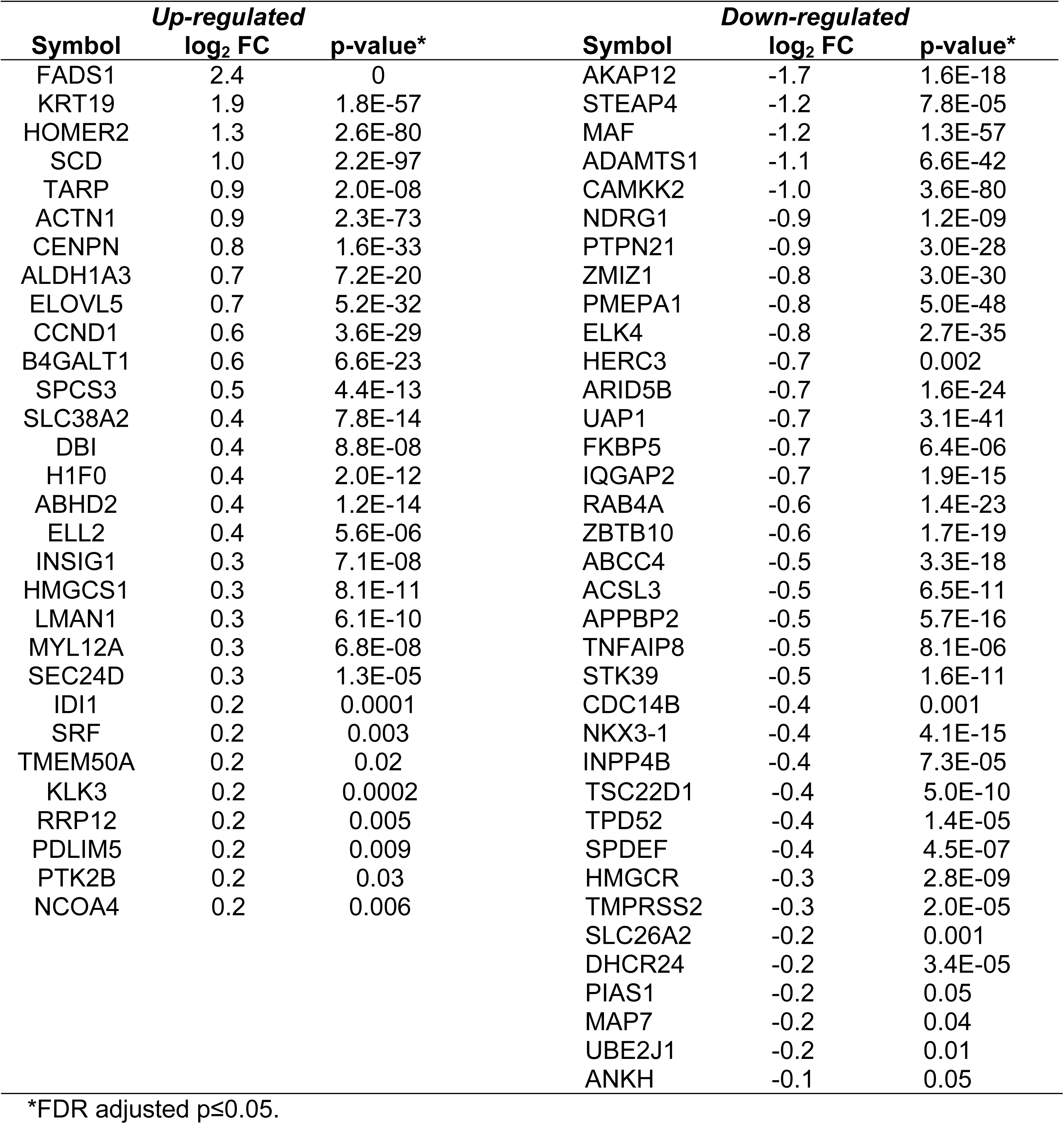
List of androgen-responsive genes differentially regulated by ligand-activated ERβ in prostate cancer cells.

Table 3: Known AR regulated genes up or down-regulated in LNCaP cells expressing ERβ and treated with R1881 and LY3201 compared to control LNCaP cells treated with R1881 and LY3201.

### ERβ reduces activation of the p (ARR) _2_ PB-LUC reporter in LNCaP and 22Rv1 cells and reduces AMPK phosphorylation

We used AR-reporter constructs to investigate whether AR activity is directly affected by ERβ in ERβ-overexpressing LNCaP cells as well as 22Rv1 PCa cells. Ligand-treatment of transfected cells showed ERβ-dependent inhibition of AR-reporter activity compared to control transfected cells (see Figure 2 E and D for luciferase assay).

### ERβ-regulation of CAMKK2 and its downstream targets

Another AR-regulated gene, CAMKK2, which has been shown previously to affect PCa survival, metabolism, cell growth, and migration[4], was repressed by ligand-activated ERβ (see Figure 3A). To determine whether events downstream of CAMKK2 were affected by the presence of activated ERβ, we analyzed the activity of AMPK by measuring its phosphorylated form (pAMPK). There was a clear reduction in pAMPK following treatment with LY3201 (Figure 3 B). We also found the CAMKK2 regulated gene TBC1D4 to be down-regulated by ERβ (Figure 3C and D). This factor is important for translocation of GLUT12 to the plasma membrane and subsequent glucose transport and has previously been shown to be up-regulated by AR [15].

**Figure 3.**
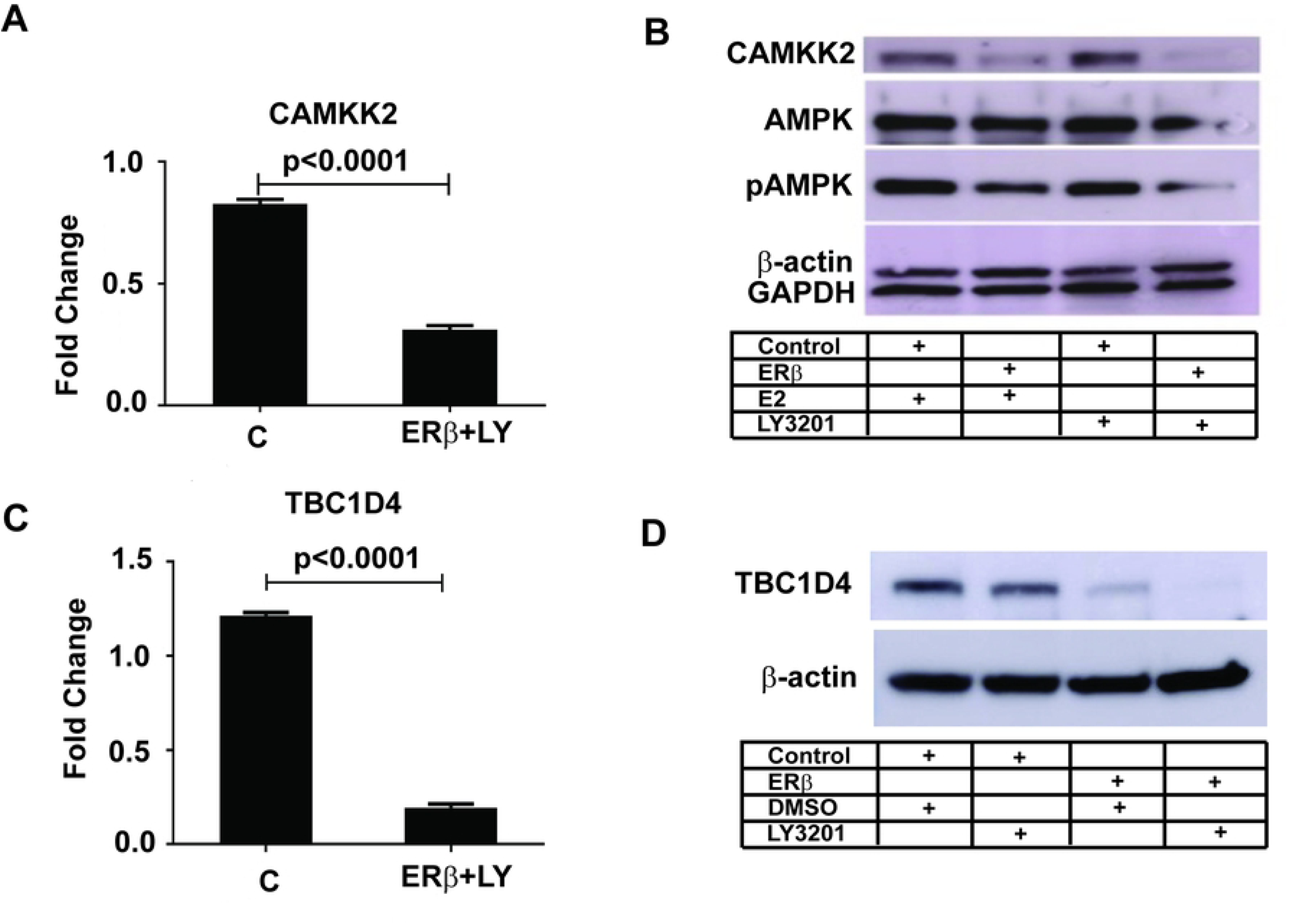
Expression and activity of AR target gene CAMKK2 are disrupted by ERβ. A, AR target gene CAMKK2 transcript levels were reduced by ERβ activation. B, CAMKK2 protein expression and activity (measured by anti-AMPK and anti-phospho-AMPK antibodies raised against the CAMKK2 substrate) were down-regulated by ERβ. C, The AR and CAMKK2 target gene TBC1D4 transcript and protein levels were reduced by activation of ERβ.

## Discussion

In the prostate of ERβ knockout mice, there is an increase in the number of basal cells (p63-positive) and poorly differentiated intermediary cells, [10] as well as a decrease in fully differentiated luminal cells[11, 12]. ERβ is expressed in both luminal (AR-positive) and basal (AR-negative) cells. From previous studies we have found that ERβ has an anti-proliferative effect in PCa by down regulating Skp2 and up-regulating p27 KIP1 protein.

In the present study, we specifically examined the functions and potential mechanisms of action of ERβ in AR-positive LNCaP cells by RNA-seq. Consistent with its function as a ligand-dependent transcription factor, overexpression of ERβ elicited relatively little changes in the LNCaP transcriptome in a ligand-independent fashion, but treatments with ER ligands E2 or ERβ-specific ligand LY3201 resulted in altered expression of thousands of genes. As expected, GSEA results revealed significant enrichment of ER-regulated genes among those up-regulated by ligand treatment. Of particular interest is the down-regulation of genes involved in androgen response. GSEA results and follow-up studies on the expression and activity of AR and target genes shown herein provide evidence that the tumor suppressor actions of ERβ in PCa are due to its negative regulation of AR signaling. AR expression and/or activity is often increased in primary PCa, as well as in metastatic PCa. In the present study we found that in LNCAP cells overexpressing ERβ, AR activity is repressed by ERβ-ligands, as were the classical AR targets c-Myc, FKBP5 and CAMKK2. TBC1D4 is the most down-regulated AR-responsive gene and a major regulator of glucose uptake in the prostate by inducing membrane localization of Glut12. As TBC1D4 is a down-stream target of both CAMKK2 and AR, there is indication that ERβ is capable of repressing glucose metabolism in the prostate. The two most up-regulated AR responsive genes FADS1 and KRT19 is likely to be targets of both AR and ERβ, where FADS1 have been shown to be regulated by estrogen [16], and KRT19 expression correlating to estrogen receptor expressing breast cancer [17].

The repression of AR by ERβ is not limited to LNCaP cells, but also occurs in another androgen-responsive cell line, 22Rv1, which expresses both normal AR as well as the AR variant AR7. In addition to regulating AR, GSEA and additional gene ontology analysis of genes differentially expressed following ERβ activation revealed an enrichment of genes involved in cancer-related processes, including apoptosis, response to hypoxia, KRAS signaling, and key metabolic pathways. Another pathway that can drive prostate cancer is CYP epoxygenases which catalyze the formation of epoxyeicosatrienoic acids (EETs) from arachidonic acid. The CYPS involved belong to family 4A, as well as 2U2 and 2J2 [18, 19]. CYP2U2 catalyzes conversion of arachidonic acid into two bioactive compounds, the 19- and 20-HETE. Fatty acid epoxides are short-lived because they are hydrolyzed to less active or inactive dihydroxy-eicosatrienoic acids by soluble epoxide hydrolases [20]. Thus the activity of these epoxides in also dependent upon the expression levels of epoxide hydroxylases.

Yet another pathway factor associated with increased risk of PCa is reduction in vitamin D. The CYP involved in the first step in the activation of vitamin, is the 25-hydroxylase (CYP 2R1). In addition to being the precursor of the active hormone, 1, 25(OH)_2_D3, 25hydroxy vitamin D has actions of its own in the prostate. It is involved in keeping metabolism in the prostate in the normal mode i.e., oxidative phosphorylation predominating over glycolysis. A reduction of oxidative phosphorylation occurs when prostate cells become malignant [21].

Another interesting finding is cytochromes P-540 involved in formation of epoxides from fatty acids and the synthesis of 25-hydroxy vitamin D was reduced to 50 % and the lysophosphatidic acid receptor LPAR3 to 25 % of levels in untreated cells. These findings reveal that even in malignant cells, introduction of ERβ down regulates AR signaling as well as other possible drivers of PCa, fatty acid epoxygenases, (lysophosphatidic/ GPR pathway) and vitamin D synthesis. Whether the effects of ERβ on these genes are the consequences of its interaction with AR or through independent mechanisms remain to be determined. Nonetheless, these findings reveal that re-expression and activation of ERβ can suppress oncogenic mechanisms in androgen-responsive cancer cells. Nonetheless, these findings reveal that re-expression and activation of ERβ can suppress oncogenic mechanisms in androgen-responsive cancer cells. These findings reveal possible early therapeutic interventions in androgen-responsive PCa through activation of ERβ. Since expression of these potential drivers are high even in the absence of ERβ signaling, they could be targeted directly for treatment of PCa.

## Acknowledgements

This study was supported by grants from the Brockman Foundation (000176005) and the Robert A Welch Foundation (E-0004)

